# Transcriptional analysis of developing *Aspergillus fumigatus* biofilms reveals metabolic shifts required for biofilm maintenance

**DOI:** 10.1101/2025.06.02.657448

**Authors:** Charles Puerner, Kaesi A. Morelli, Joshua D. Kerkaert, Jane T. Jones, Katherine G. Quinn, Sandeep Vellanki, Robert A. Cramer

**Author notes:** Corresponding Author: Robert A. Cramer, Ph.D. CP, KAM, and JDK contributed equally to this study.

## Abstract

*Aspergillus fumigatus* is a filamentous fungus found in compost and soil that can cause invasive and/or chronic disease in a broad spectrum of individuals. Diagnosis and treatment of aspergillosis often occur during stages of infection when *A. fumigatus* has formed dense networks of hyphae within the lung. These dense hyphal networks are multicellular, encased in a layer of extracellular matrix, and have reduced susceptibility to contemporary antifungal drugs, characteristics which are defining features of a microbial biofilm. A mode of growth similar to these dense hyphal networks observed *in vivo* can be recapitulated *in vitro* using a static, submerged biofilm culture model. The mechanisms underlying filamentous fungal cell physiology at different stages of biofilm development remain to be defined. Here, we utilized an RNA sequencing approach to evaluate changes in transcript levels during *A. fumigatus* biofilm development. These analyses revealed an increase in transcripts associated with fermentation and a concomitant decrease in oxidative phosphorylation related transcripts. Further investigation revealed that ethanol and butanediol fermentation is important for mature biofilm biomass maintenance. Correspondingly, a gene (silG), a predicted transcription factor, was observed to also be required for mature biofilm biomass maintenance. Taken together, these data suggest temporal changes in *A. fumigatus* metabolism during biofilm development are required to maintain a fully mature biofilm.

**IMPORTANCE:** *Aspergillus fumigatus* is the most common etiological agent of a collection of diseases termed aspergillosis. Invasive <u>P</u>ulmonary <u>A</u>spergillosis (IPA), a severe form of aspergillosis, is highlighted by invasive growth of fungal hyphae into host lung tissue. Strains that are susceptible to antifungal therapies *in vitro* frequently fail to respond to treatment *in* vivo, resulting in high mortality rates even with treatment. It is now appreciated that this decreased antifungal efficacy *in vivo* is, in part, likely due to biofilm-like growth of the fungus. *A. fumigatus* biofilms have been shown to develop regions of limited oxygen availability that are hypothesized to induce cell quiescence and drug resistance. Understanding the mechanisms by which *A. fumigatus* induces, develops, and maintains biofilms to evade antifungal therapies is expected to illuminate biofilm-specific therapeutic targets. Here we present transcriptomics data of developing *A. fumigatus* biofilms and from these data define genes related to fungal fermentation and regulation of transcription important for maintenance of mature *A. fumigatus* biofilms.

## INTRODUCTION

*Aspergillus fumigatus* is a ubiquitous filamentous fungus that causes a broad spectrum of acute and chronic diseases including <u>IPA</u>. Mortality of IPA remains high, even with antifungal treatment, and strains that respond to antifungal drugs *in vitro* frequently fail to respond to corresponding treatment *in vivo* (Perea & Patterson, 2002). As diagnosis of aspergillosis can be challenging, antifungal treatment of IPA typically does not occur until the fungus has had sufficient time to establish mature, dense hyphal networks within lung tissue. These hyphal networks at sites of infection have hallmark characteristics of a microbial biofilm, including multicellularity, extracellular polysaccharide secretion, surface adherence, and limited oxygen availability (Beauvais et al., 2014; Kowalski et al., 2020; Loussert et al., 2010; Mowat et al., 2007; Müller et al., 2011; Rajendran et al., 2011).

*A. fumigatus* biofilms have been shown to generate regions of limited oxygen availability both *in vitro* and *in vivo* (Grahl et al., 2011; Kowalski et al., 2020). *In vitro*, this low oxygen environment is in part responsible for the observed increasing antifungal resistance as biofilms develop (Kowalski et al., 2020). However, the mechanisms through which *A. fumigatus* biofilms fully develop and maintain their antifungal drug resistance remain ill-defined (Liu et al., 2022; Morelli et al., 2021). One proposed mechanism is through the presence of extracellular DNA in the extracellular matrix (Rajendran et al., 2013). Identifying biofilm-specific antifungal drug resistance mechanisms is expected to help develop new therapeutic targets to increase antifungal efficacy *in vivo* when robust biofilm-mediated infections are established.

To help fill the knowledge gap on *A. fumigatus* biofilm development mechanisms, we utilized a submerged biofilm culture model to identify genes expressed at different timepoints of biofilm development. We focus on pathways that are important in the establishment and maintenance of mature biofilms as well as pathways that are important for mediating fungal metabolism at different stages of biofilm development. The transcriptomics data suggest a shift from oxidative phosphorylation-mediated energy generation towards a more fermentative metabolism as the biofilm matures. In support of this observation, we observed that loss of genes required for ethanol and butanediol fermentation reduced the biomass of mature biofilms. Additionally, we identified a previously uncharacterized predicted transcription factor whose transcript levels increase throughout biofilm development and is consequently required for maintenance of biofilm biomass. These data provide insight into the genes and pathways important for *A. fumigatus* submerged biofilm development and reveal a previously unidentified transcriptional regulator as a mediator of mature biofilm maintenance.

## RESULTS

### Biofilm developmental stages are transcriptionally unique

To identify genes and pathways that are important during distinct stages of *A. fumigatus* submerged biofilm development, we utilized an RNA-Seq approach. The *A. fumigatus* laboratory strain, CEA10 (also known as Fungal Genetics Stock Center strain A1163), was utilized to generate submerged biofilms in glucose minimal medium for 12-, 18-, 24-, and 30-hours of biofilm development (**Fig. 1)**. These time points were chosen based on our previous analysis of biofilm architecture and antifungal drug susceptibility in this model at these timepoints (Kowalski et al., 2020). Total RNA was extracted from replicate biofilms at each timepoint and analyzed by paired-end Illumina RNA sequencing. We acquired a mean of 10.1 M reads per sample (SD = 5.5 M) and had a mean mapping rate of 81.2% (SD = 3.6%) to the A1163 reference genome (Fedorova et al., 2008).

**Figure 1:**
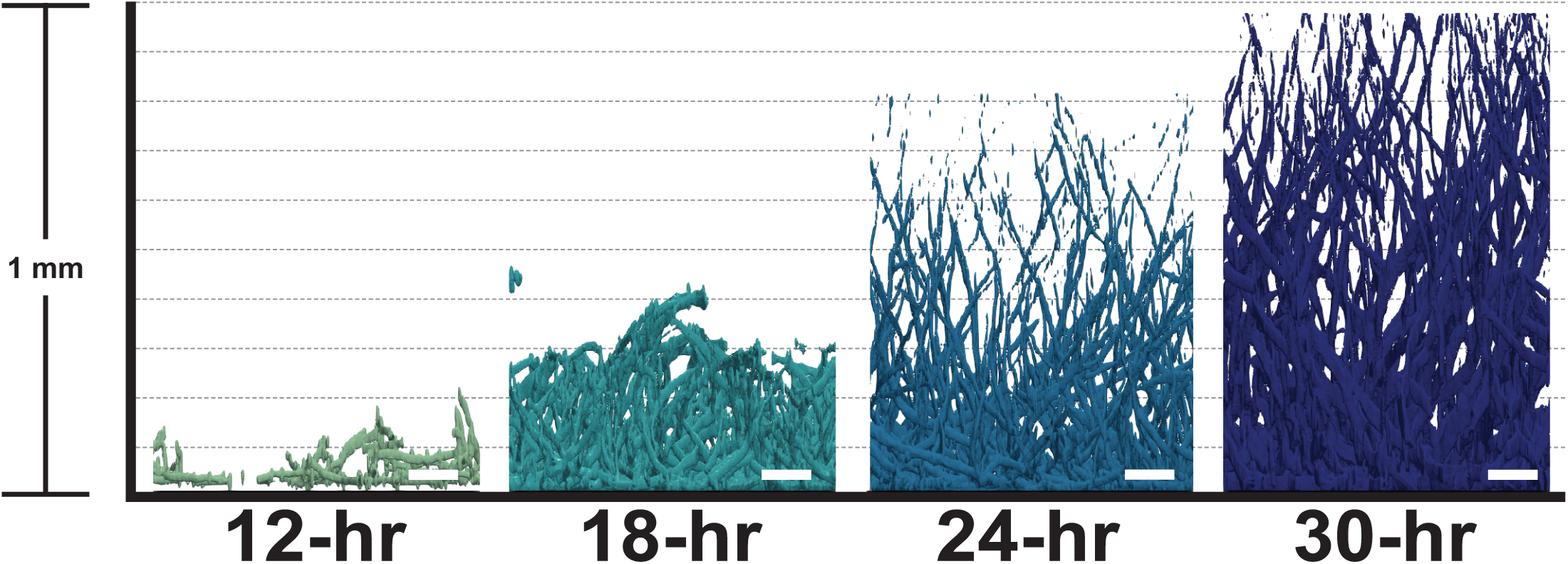
Morphology of biofilm development. Z-stack images acquired at 12-, 18-, 24-, and 30-hour timepoints during development of the *A. fumigatus* biofilm. Biofilms are stained with calcofluor white prior to imaging. Fluorescence images were rendered using BiofilmQ and Paraview software. Scale bar is 100 µm.

Exploratory analysis revealed each biofilm stage is transcriptionally unique. Global Euclidian clustering of all samples shows samples within each timepoint are closer than samples from other timepoints (**Fig. 2A**). Principal component analysis of the top 2000 most variable transcripts further confirms the inter-timepoint clustering and a clear intra-timepoint separation **(Fig 2B**). The top 2000 most variable genes were used in this analysis as this captured the majority of the variation in the dataset (**Fig. S1**). Samples within the same timepoint clustered strongly together, and principal component (PC) 1 (78.9% of variance) is explained by a clear stepwise separation of biofilm development along this PC while PC2 (13.5% of variance) is seemingly explained by developing biofilm versus immature and mature biofilms. From both exploratory analyses we determined that 12-hour biofilm samples are transcriptionally distinct from the three later biofilm stages. A hierarchical clustering analysis of transcript abundance dynamics of the top 2000 most variable transcripts revealed potential clusters of genes important for each stage of development (**Fig. 1C**). For example, clusters of genes with transcripts that are increased at 18-hour when compared to 12-hour but subsequently reduced in abundance at 24-hour compared to 12-hour, suggesting these transcripts are specifically important for the 18-hour stage of biofilm development. Similar patterns are observed at later timepoints. For example, a cluster of transcripts that are strongly increased at 30-hours compared to 24-hours but reduced or relatively unchanged in all other comparisons have been identified (**Table S1**).

**Figure 2:**
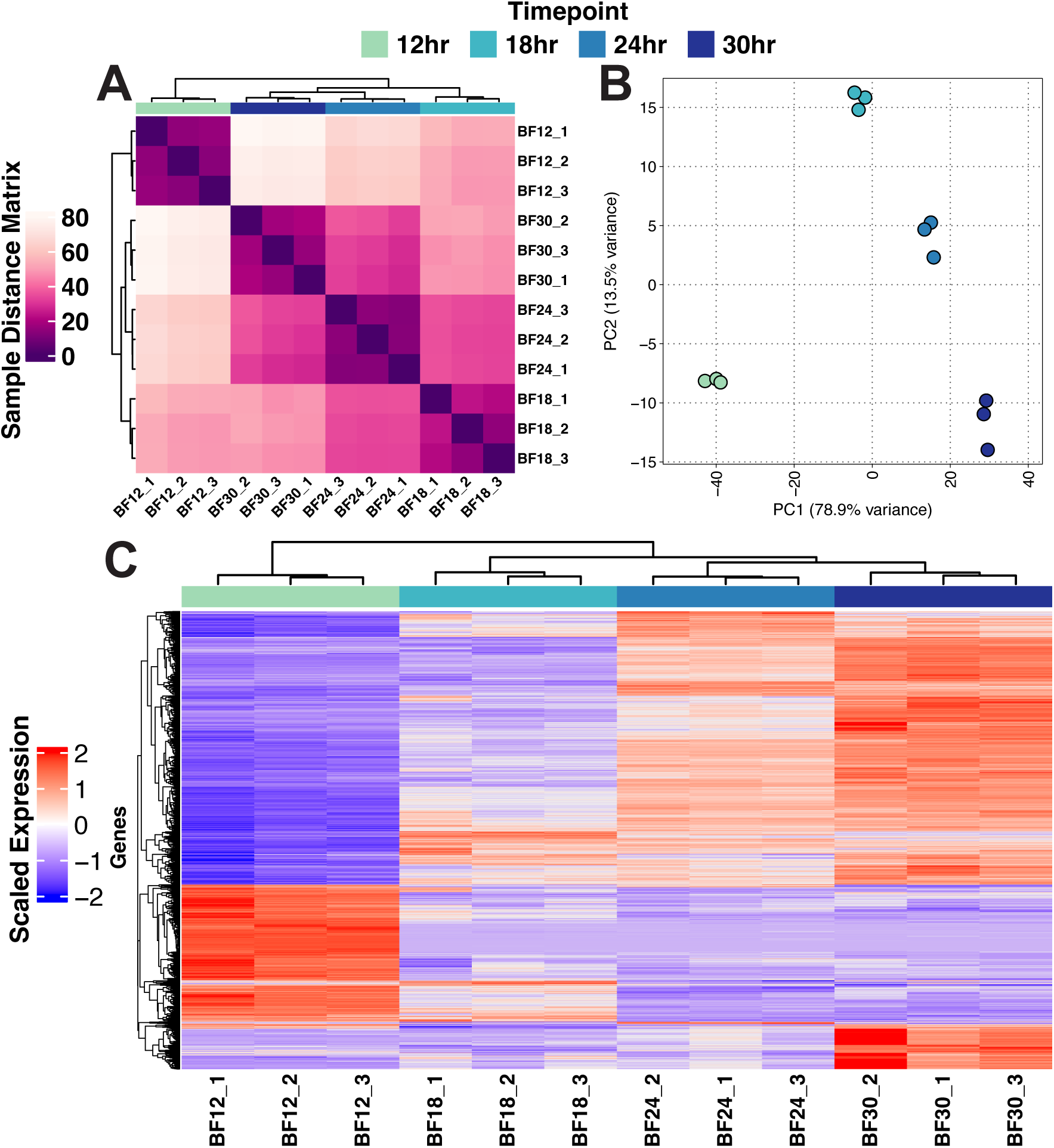
Exploratory analysis of biofilm transcriptional time course. **A**) Euclidean distance matrix representing distance relationships between samples. **B**) Principal component analysis shows separation of time course in the first principal component. **C**) Hierarchical clustering of the top 2000 most variable genes shows progression of biofilm development with the 12-hour timepoint showing a unique state compared to remaining timepoints. Scaled vst normalized values are shown in heatmap. Heatmap shows likely timepoint specific gene expression patterns.

### Investigation of timepoint specific expression patterns

We utilized a pairwise differential expression analysis comparing all the possible combinations of pairwise comparisons between the 4 timepoints (6 comparisons) (**Table 1, S2**). In order to assess the overlap in transcript abundance between timepoints we generated upset plots for transcripts that are significantly increased in abundance and transcripts that are significantly decreased in abundance (**Fig. 3**). There was a strong overlap in the number of transcripts that are both increased and decreased in all timepoints compared to the 12-hour timepoint, further demonstrating there is an important transcriptional change in this model leading to biofilm development that occurs between the 12-hour and 18-hour timepoints. Additionally, these plots indicate the potential for stage-specific transcript regulation.

**Figure 3:**
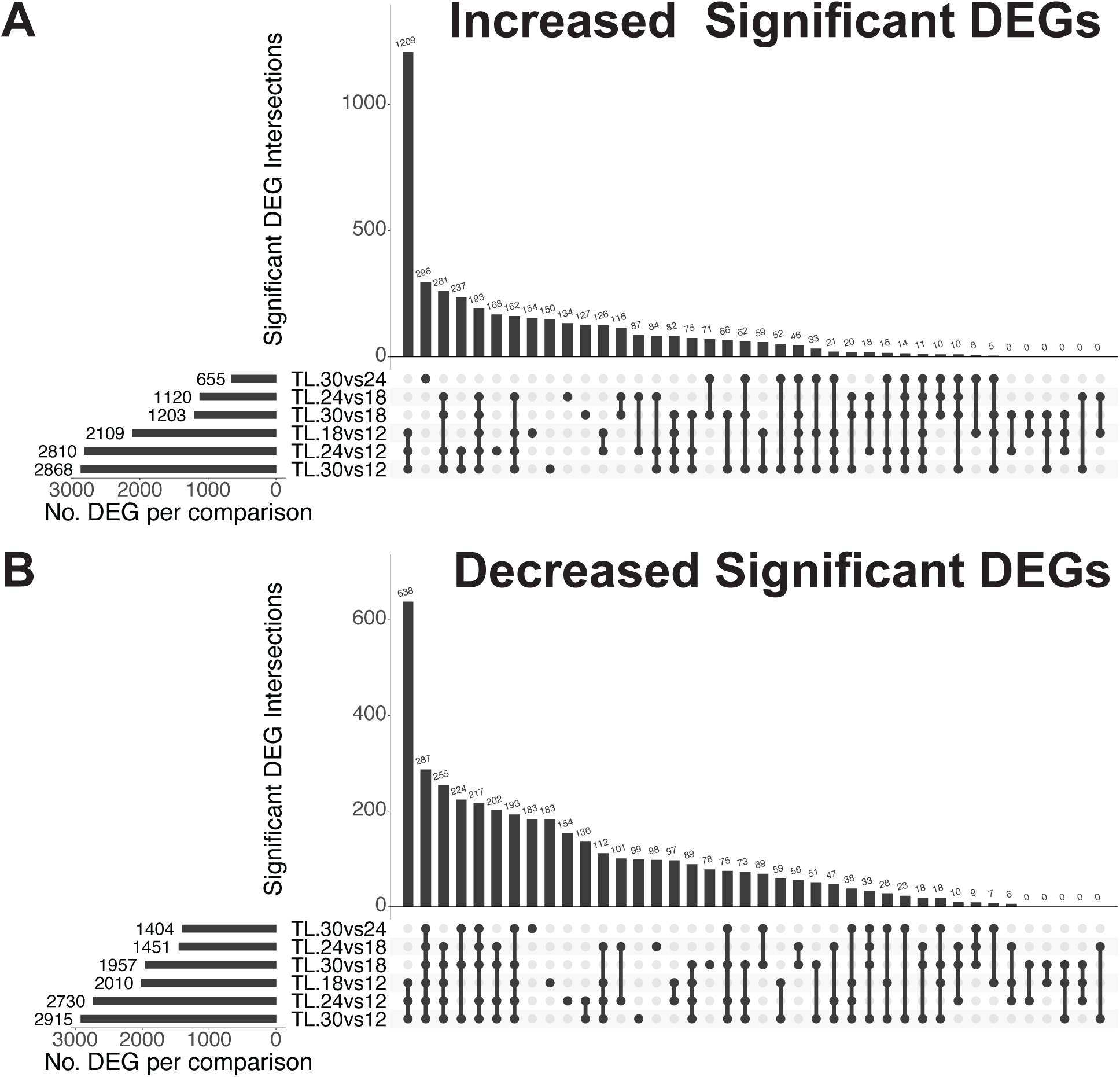
Differential expression across timepoints. Upset plots are used to investigate overlapping significant differentially expressed genes (sDEGs) in the time course analysis. Gene expression was compared pairwise across timepoints to identify sDEGs and upset plots were used to analyze overlaps in sDEGs between comparisons with **A**) showing increased transcripts and **B**) showing decreased transcripts. An adjusted p-value cutoff of 0.05 was used to identify sDEGs.

**Table 1.** Summary of differential gene expression analysis.

| Comparison | LogFC_0 total | LogFC_0 Increased | LogFC_0 Decreased | LogFC_1 total | LogFC_1 Increased | LogFC_1 Decreased | LogFC_2 total | LogFC_2 Increased | LogFC_2 Decreased |
| --- | --- | --- | --- | --- | --- | --- | --- | --- | --- |
| BF18hr_vs_BF12hr | 4119 | 2010 | 2109 | 705 | 460 | 245 | 470 | 271 | 199 |
| BF24hr_vs_BF12hr | 5540 | 2730 | 2810 | 1153 | 859 | 294 | 787 | 567 | 220 |
| BF30hr_vs_BF12hr | 5783 | 2915 | 2868 | 1456 | 1162 | 294 | 1062 | 838 | 224 |
| BF24hr_vs_BF18hr | 2571 | 1451 | 1120 | 279 | 259 | 20 | 159 | 150 | 9 |
| BF30hr_vs_BF18hr | 3160 | 1957 | 1203 | 687 | 651 | 36 | 454 | 439 | 15 |
| BF30hr_vs_BF24hr | 2059 | 1404 | 655 | 234 | 224 | 10 | 153 | 151 | 2 |

Through the exploratory and differential expression analysis we observed clusters of transcripts that are likely regulated at specific timepoints. To explore this observation further, an analysis to identify genes that have transcript levels regulated at specific timepoints was conducted using four criteria: 1) significantly increase in transcript abundance and remain increased for all subsequent timepoints (plateau), 2) significantly decrease in abundance and remain low for all subsequent timepoints (lowland), 3) significantly increase in abundance at a specific timepoint (peak), and 4) significantly decrease in abundance at a specific timepoint (valley). The analysis was conducted using log_2_ fold-change and adjusted p-value cutoff of significance with criteria used for each timepoint outlined in **table 2**. From this analysis we identified clusters of plateau, lowland, peak and valley transcripts for each timepoint (**Fig 4, table S3**). We utilized an enrichment analysis using the FunCat categories related to metabolism from the FungiFun3 resource (Garcia Lopez et al., 2024) (**table S4**).

**Figure 4:**
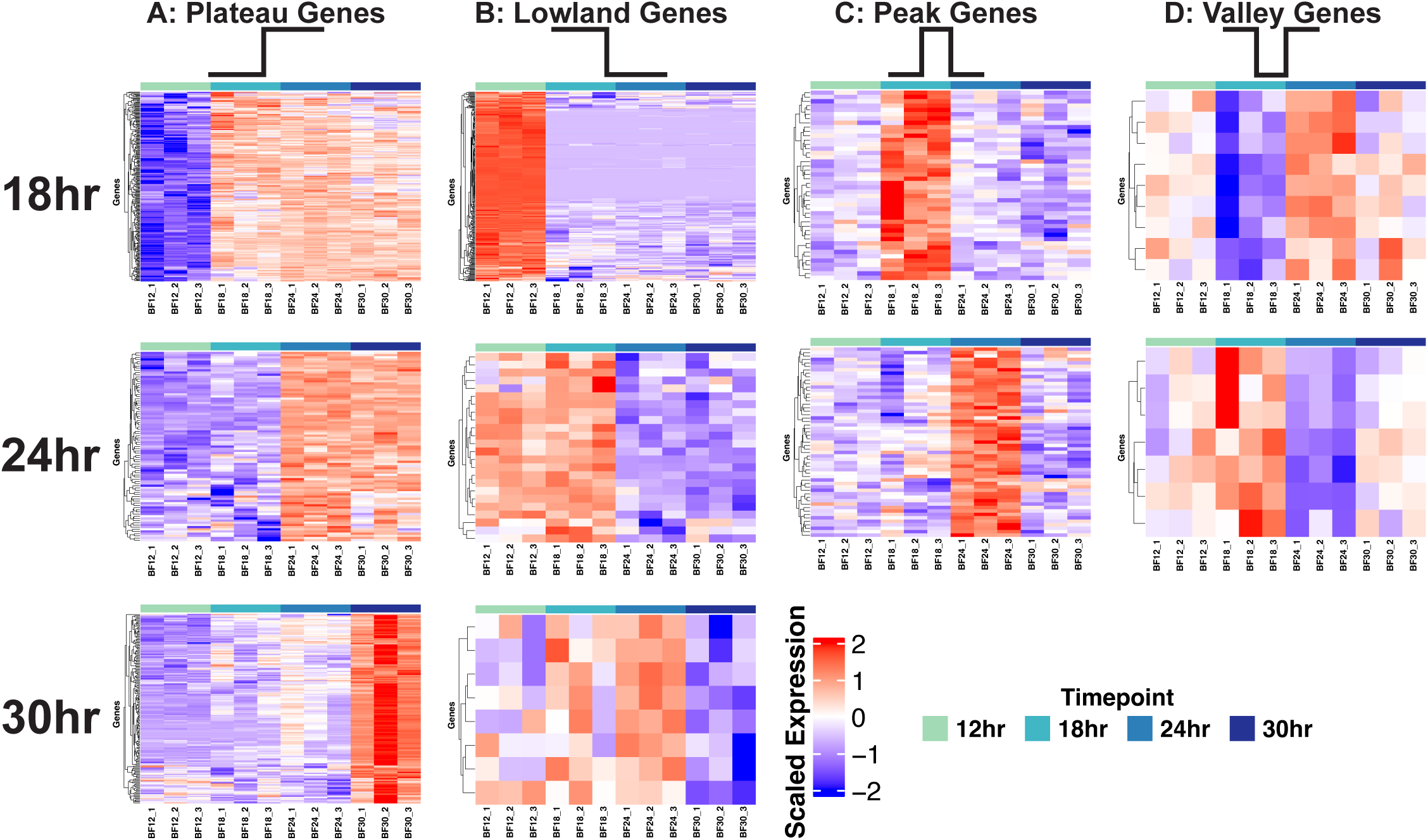
Identification of dynamic gene expression patterns. Using differential expression data (log2FC and adjusted p-value) dynamic gene expression patterns were identified for each timepoint. **A**) Plateau genes increase in expression at given timepoint and remain increased during development. **B**) Lowland genes decrease in expression at a given timepoint and remain lowly expressed during the remainder of development. **C**) Peak genes increase in expression specifically at 18- or 24-hour timepoints. **D**) Valley genes decrease specifically at 18- or 24-hour timepoints. Scaled CPM values are shown.

**Table 2.**
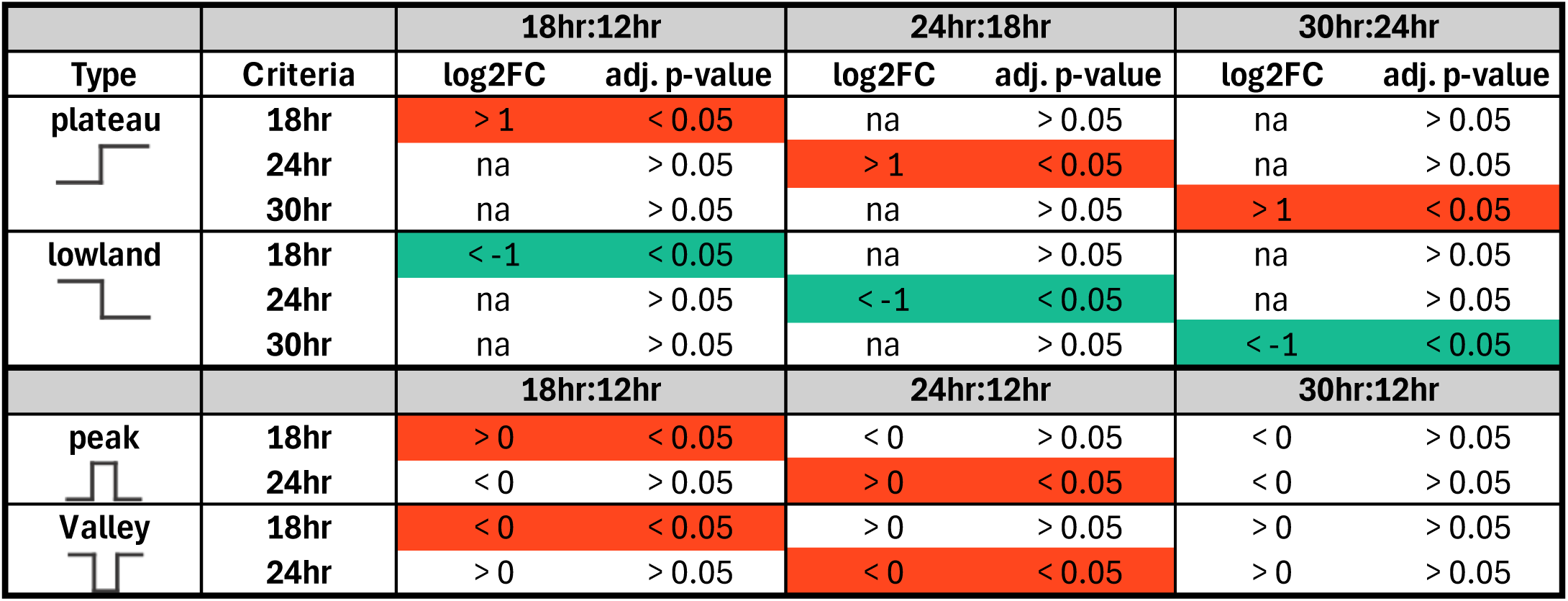
Criteria used to identify peak, plateau, and lowland genes.

From this analysis it is suggested that metabolic shifts take place over the course of biofilm development. For example, Funcat categories for secondary metabolism and toxins were significantly enriched in the 24- and 30-hour plateau gene lists. Due to this finding, we investigated the transcript abundances of several secondary metabolite and related pathways identified using the KEGG database (**Fig. S2**). We observed several biosynthetic gene clusters for secondary metabolite biosynthesis with increased transcript abundance in late stages of biofilm development (**Fig. S3)**. Specifically, we found the following biosynthetic pathways to have increased transcripts: Fumagillin, Fumigaclavine, Fumiquinazoline, and Fumitremorgin. As these pathways appear to be turning on as the biofilms mature, it is an intriguing possibility that they are involved in biofilm maintenance.

### Biofilms switch from respiratory metabolism to fermentative metabolism

To further define the metabolic changes occurring during biofilm development, we compared transcript levels of genes associated with oxidative phosphorylation and fermentative metabolism at each stage of biofilm development. Using the KEGG database we identified genes associated with oxidative phosphorylation to subset our data (Kanehisa & Goto, 2000). In support of our hypothesis that metabolic re-wiring occurs during biofilm development, transcripts for the majority of genes associated with oxidative phosphorylation were highest at the 12-hr timepoint compared to all other timepoints **(Fig. 5A).** Biofilms at 18 hours also had relatively high transcript abundance of genes encoding proteins predicted to function in oxidative phosphorylation, though some heterogeneity is observed at this timepoint. By the 24-hour timepoint, there is a relative reduction in the mRNA abundance of oxidative phosphorylation related genes. Oxidative phosphorylation-associated transcripts are further reduced in 30-hour biofilms.

**Figure 5:**
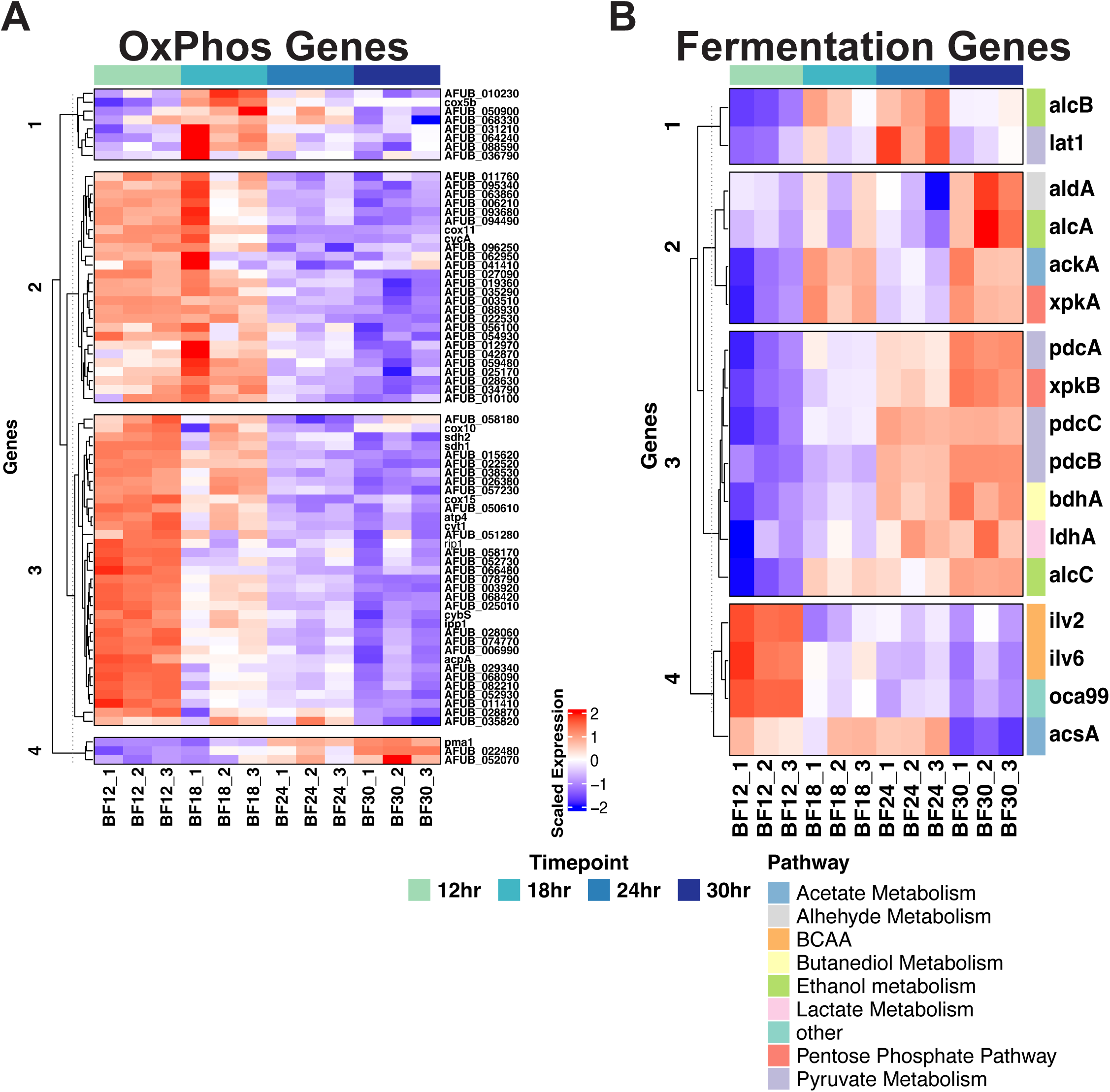
Gene expression of oxygen dependent and independent energy production pathways. Gene expression from genes within (**A**) oxidative phosphorylation (OxPhos) and (**B**) fermentation pathways found a general decrease in oxidative phosphorylation genes and a general increase in fermentative genes. Scaled CPM values are shown in the heatmaps.

Conversely, comparison of transcript levels of genes encoding proteins associated with fermentative metabolism and the phosphoketolase pathway revealed an opposite transcript abundance pattern compared to the oxidative phosphorylation related genes **(Fig. 5B)**. Transcripts from genes involved in fermentation and the phosphoketolase pathway were relatively reduced in 12- and 18-hour biofilms while being relatively more abundant in late-stage biofilms. Specifically, the 24- to 30-hour timepoints biofilms have increased mRNA abundance for genes involved in ethanol, butanediol, and lactate fermentation pathways and the phosphoketolase pathway. Interestingly our data indicates that after 12 hours there is an apparent decrease in transcripts associated with branched chain amino acid metabolism pathways. These data suggest that mature biofilms enter a state of reduced oxidative phosphorylation and potentially rely on fermentative metabolism for redox homeostasis during biofilm maintenance.

### Fermentation pathways are important for biofilm development

Transcriptionally, we observed that transcripts encoding proteins related to a metabolic shift from respiration at early timepoints to fermentation at later timepoints is likely important for biofilm maturation. We subsequently tested the hypothesis that fermentation pathways are important for biofilm maturation and/or maintenance through generation of null mutants of genes involved in two fermentative pathways, ethanol and butanediol metabolism.

Our group previously investigated the alcohol dehydrogenase involved in ethanol production, AlcC, observing that loss of *alcC* resulted in a reduction in fungal burden in a murine model of invasive pulmonary aspergillosis (Grahl et al., 2011). The cause for the reduction in fungal burden of the Δ*alcC* strain remains ill-defined and the role of AlcC in biofilm formation was not explored. One possibility given *alcC* transcript dynamics in the submerged biofilm model is that loss of *alcC* impacts the ability of the fungus to form or maintain a mature biofilm. To test this hypothesis, we quantified biofilm biomass of the Δ*alcC* strain at early (18hr) and late (40hr) biofilm timepoints (**Fig. 6A,D**). We observed that the CEA10 Δ*alcC* strain has a slight however not statistically significant reduction in overall biomass at a late stage of biofilm maintenance. We chose to utilize an additional late-stage biofilm at 40hr as we were interested in how the observed transcriptional change at 30hr would impact biofilm maintenance.

**Figure 6.**
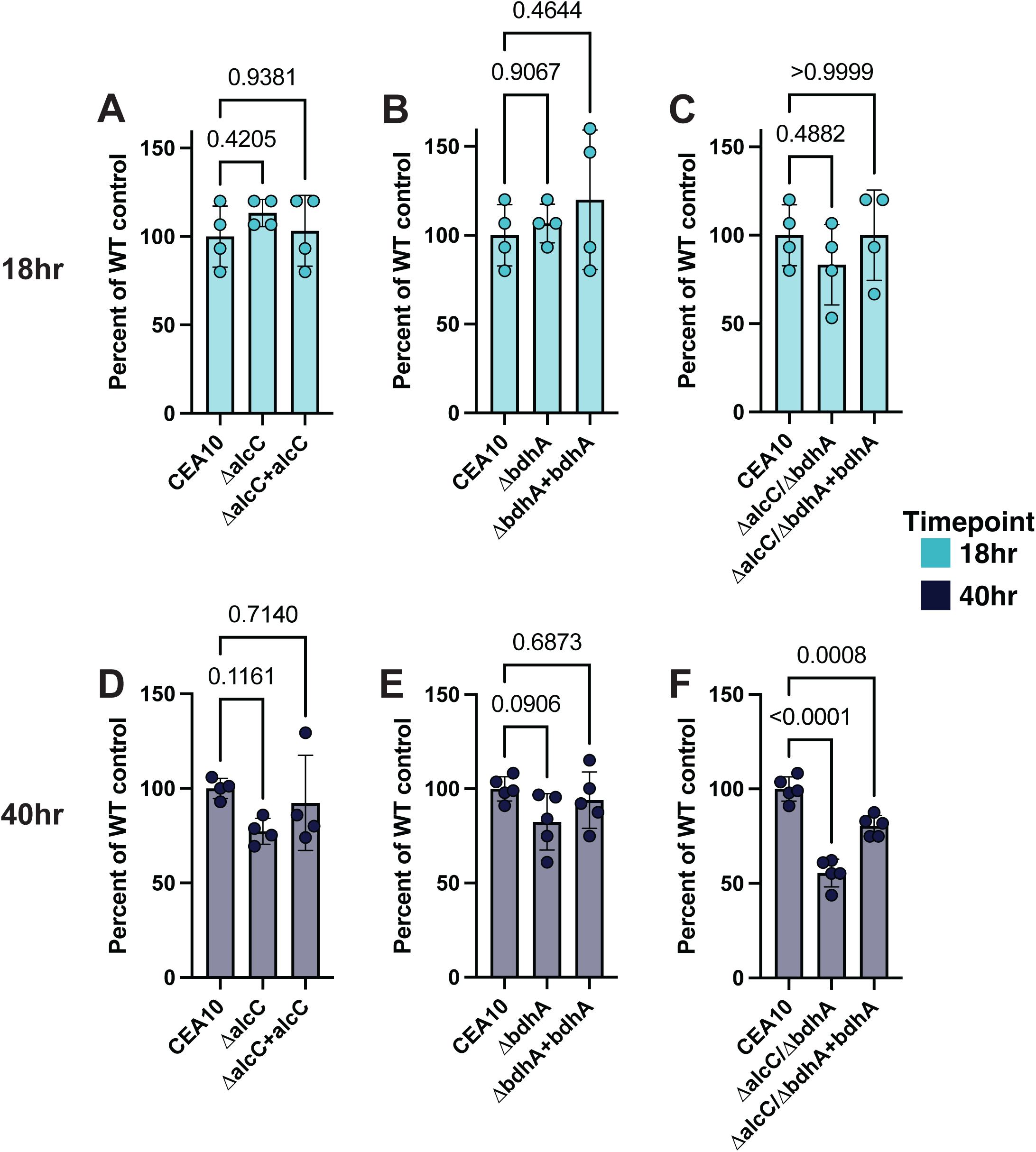
Fermentation metabolism is important for biofilm maturation. **A-C**) Biofilm biomass of fermentation mutants at 18-hours of growth. **D-F**) Biofilm biomass of fermentation mutants at 40-hours of growth. Points indicate biological replates (averages of two technical replicates) with an n=4-5 biological replicates. Statistics analysis is an ordinary one-way ANOVA with a Dunnett’s multiple comparison test.

Given that our transcriptional analysis indicates several putative fermentation pathways are induced in the biofilm at later stages, it is possible that loss of a single fermentation pathway in our model does not substantially impact biofilm maintenance. We hypothesized that removing one pathway would shift the metabolic flux to other fermentative pathways. In support of our hypothesis, we observed that the Δ*alcC* mutant has an increased production of acetoin, an intermediate product of the butanediol fermentation pathway, when shaking batch cultures grown in atmospheric oxygen conditions are shifted to low oxygen conditions for 48 additional hours (**Fig. S4 A,D)**. Despite the transcript profile suggesting an increase in butanediol production in our submerged biofilm model, we were unable to detect acetoin from the submerged biofilm cultures (**data not shown**). It is possible that the low culture biomass in our model precludes detection of fermentation products in the culture medium. We consequently impaired the butanediol fermentation pathway by generation of a null mutant for the gene encoding the putative butanediol dehydrogenase AFUB_031610, herein named *bdhA*. We identified this gene and encoded amino acid sequence in the *A. fumigatus* genome as the top hit in a reciprocal blast search for the *Saccharomyces cerevisiae* Bdh1p sequence. An EMBOSS Needle alignment between AFUB_031610 and Bdh1p revealed 36.4% identity and 56.7% similarity (Madeira et al., 2024). Similar to the Δ*alcC* strain, the Δ*bdhA* strain has a slight but not statistically significant defect in biofilm formation indicated by a reduction in biomass at a 40-hour submerged biofilm timepoint (**Fig. 6B,E**). We did not detect acetoin accumulation in the Δ*bdhA* strain, providing support that the ethanol production pathway is the preferred *A. fumigatus* fermentation pathway (**Fig. S4B,D**). We next impaired both the ethanol and butanediol fermentation pathways by generating a double null mutant strain, Δ*bdhA/ΔalcC.* We tested this strain for biofilm formation at 18- and 40-hours and observed the double mutant leads to a statistically and biologically significant ∼50% reduction in biofilm biomass compared to WT control, strongly indicating that fermentative metabolism is important for the maintenance of an *A. fumigatus* biofilm in this model (**Fig. 6C,F**). Restoration of *bdhA* expression in the double mutant restored biomass levels to that of the Δ*alcC* strain as expected. We quantified acetoin in the Δ*alcC/ΔbdhA* strain and found a similar level as the Δ*alcC* strain (**Fig. S4C,D)** suggesting other fermentative pathways derived from pyruvate, such as lactate fermentation, likely have increased flux when both ethanol and 2,3 butanediol fermentation pathways are impaired.

### The SrbA regulon is dynamically regulated during biofilm development

A likely explanation for the shift towards fermentation in late stages of biofilm development is the formation of oxygen limited environments, an established feature of *A. fumigatus* biofilms in this model (Kerkaert et al., 2022; Kowalski et al., 2019, 2020). To investigate this further we examined transcript abundance of genes regulated by SrbA, a transcription factor essential for low oxygen adaptation and biofilm development in *A. fumigatus* (**Fig. 7A**) (Chung et al., 2014; Kowalski et al., 2020; Willger et al., 2008). We found that a cluster of 32 transcripts from genes in the SrbA regulon generally increases during biofilm development. Importantly, among these increasing transcripts are the alcohol dehydrogenase *alcC* and the C-4 methyl sterol oxidase *erg25A* tracking with the general response to low oxygen. Conversely, an additional cluster of 29 transcripts generally decrease in abundance during biofilm development. Interestingly, among these transcripts are genes associated with oxygen dependent steps in ergosterol biosynthesis such as *erg1, erg3A, erg5,* and *cyp51A/B* (Jordá & Puig, 2020; Kowalski et al., 2020). In general transcripts for genes within the ergosterol biosynthesis pathways decreased as biofilms matured (**Fig. S5)**. These transcripts were previously observed to be induced in acute responses to low oxygen environments and highlights that the heterogenous biofilm microenvironment is more complex than an acute low oxygen environment (Chung et al., 2014; Kowalski et al., 2019; Losada et al., 2014). Interestingly despite the transcriptional dynamics of the SrbA regulon the transcript abundance of *srbA* remains relatively unchanged during biofilm development (**Fig. 7B**). As the transcript abundance of SrbA does not vary significantly throughout biofilm development, the changes seen in the SrbA regulon are likely due to previously described post-translational regulation of SrbA, and/or alterations in SrbA co-factors that dictate gene regulation specificity (Willger et al., 2012). Consistent with its role as a regulator of the low oxygen response, and the development of low oxygen microenvironments within the *A. fumigatus* biofilm, loss of *srbA* prevents biofilm maturation (Kowalski et al., 2020).

**Figure 7:**
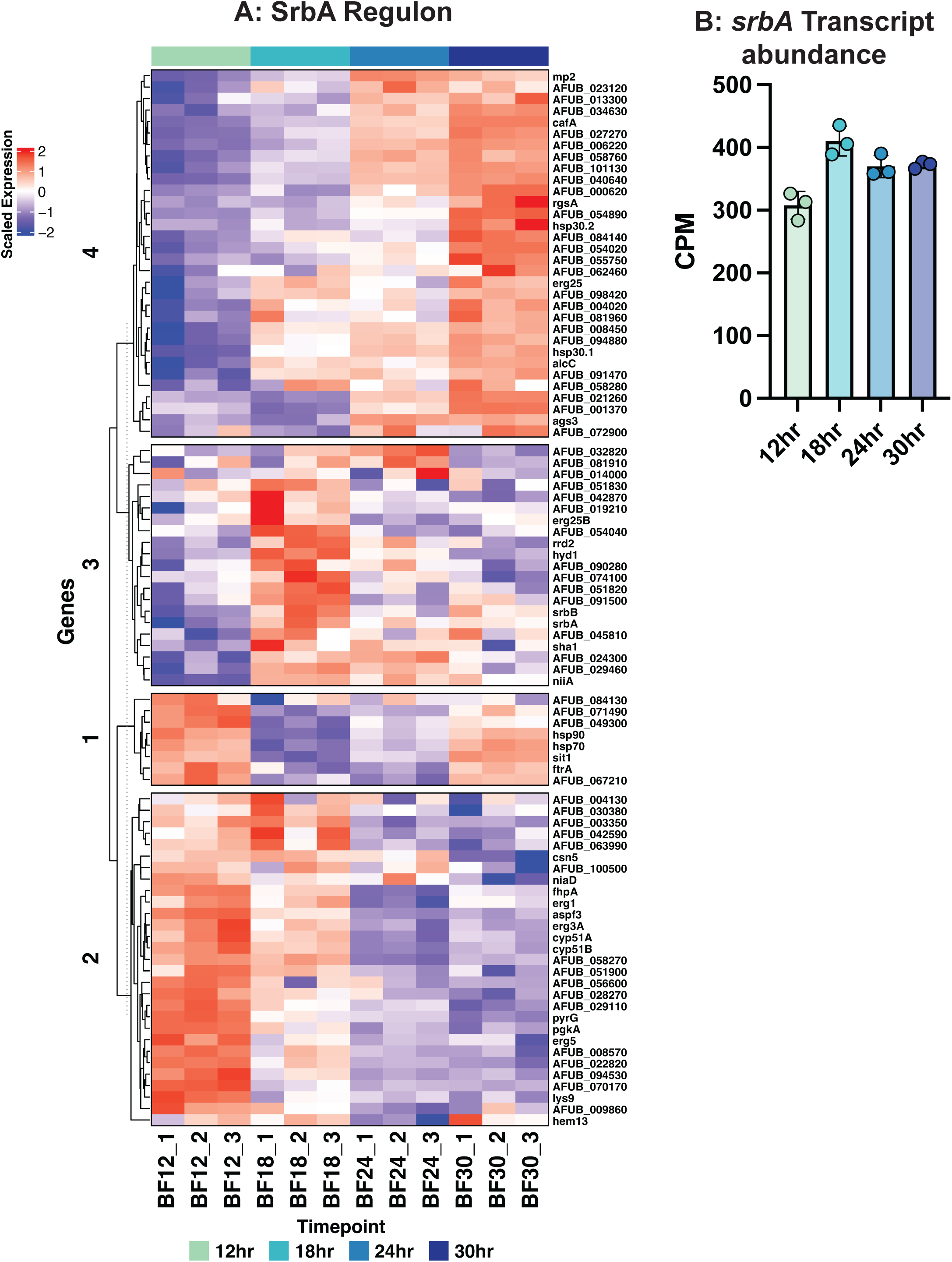
SrbA regulon activity genes during biofilm development. **A**) The expression patterns of genes within the SrbA regulon were examined using a heatmap with hierarchical clustering and 4 k-means clusters. Scaled gene expression is shown. **B**) The abundance of srbA transcripts are relatively stable during biofilm development. Graph shows CPM values of srbA transcripts at the four stages of development investigated.

### Transcription factor transcript abundance is dynamic during biofilm development

We next were interested in whether genes predicted to encode transcription factors and kinases were dynamically regulated in the developing submerged biofilm model. We subset our data for the 429 transcription factor and 108 kinase genes found in the COFUN transcription factor null mutant collection and utilized hierarchical clustering of the transcript abundances of these genes over biofilm development (Furukawa et al., 2020; van Rhijn et al., 2024). From this we found ten K-means clusters that sufficiently separate out the transcription factors and 4 clusters that separate out the kinases based on the similarity of their expression profiles. (**Fig. 8A, S6**). We chose to focus on the transcription factors in order to explore direct regulators of gene expression in this dataset. We observed several clusters of transcription factors that have dynamic transcript abundance regulation during biofilm development. For example, clusters 7, 8, and 9 have low transcript abundance in early timepoints and high transcript abundance in later timepoints while clusters 1, 2, and 3 have high transcript abundance in early timepoints and low transcript abundance in later timepoints. Transcription factors with known roles in fungal metabolism regulation showed dynamic abundance. For example, the gene *creA*, which encodes the carbon catabolite repression system transcription factor CreA (cluster 2), has decreased transcript abundance in later timepoints potentially indicating a loss of repressing carbon source(s) in late-stage biofilms (**Fig. 8B**) (Beattie et al., 2017). Additionally, we observed altered transcript abundance for metabolism regulating transcription factors *acuK* (cluster 8) and *acuM* (cluster 5) (**Fig. 8C,D)**.

**Figure 8:**
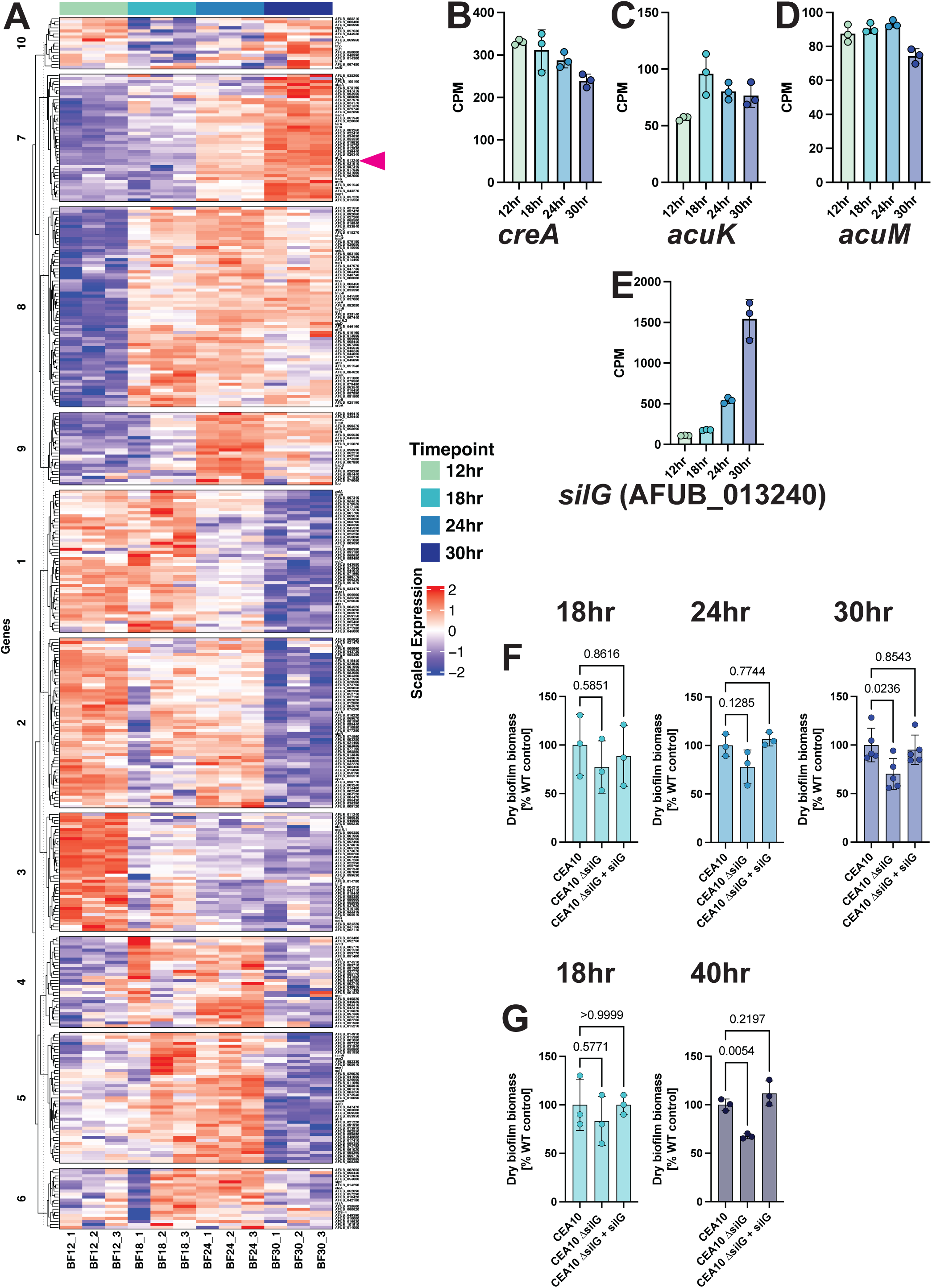
Transcript abundance of transcription factor genes is dynamic and involved in biofilm development. **A**) Heatmap showing transcript abundance of 429 transcription factors with hierarchical clustering and 10 k-means clusters. **B-D)** Transcript abundance of the metabolic transcriptional regulators *creA, acuK,* and *akuM* **E**) Transcript abundance of the transcription factor *silG* (AFUB_013240) during development (also indicated by pink arrowhead in heatmap). **F**) The Δ*silG* null mutant is impaired in biofilm development at later stages of development. Biofilm biomass of the ΔAFUB_013240 mutant at 18-, 24-, and 30-hour timepoints. **G)** The ΔsilG mutant has a further reduction in biofilm biomass production at 40-hours of growth. Dots represent biological replicates which are averages of two technical replicates with n = 3-4 biological replicates. Statistics shown are a one-way ANOVA with a Dunnett’s multiple comparison test.

### A biofilm development responsive transcription factor

Through our analysis of transcription factor transcript dynamics, we identified a potential regulator of biofilm development and maintenance, an uncharacterized putative transcription factor, AFUB_013240. AFUB_013240 transcript levels are significantly increased throughout the course of biofilm development, with its highest transcript levels observed in 30-hour biofilms **(Fig. 8E)**. Furthermore, AFUB_013240 was also observed to be significantly induced in response to low oxygen conditions in two previous studies (Kowalski et al., 2019; Losada et al., 2014). A reciprocal protein BLAST search for orthologs in *S. cerevisiae* revealed the closest yeast homolog to AFUB_013240 is the proteosome activator RPN4 (YDL020) (Xie & Varshavsky, 2001; Yau et al., 2023). However, sequence alignment using the EMBOSS Needle algorithm results in only 16.5% identity and 27.8% similarity between the *A. fumigatus* protein and yeast RPN4p, largely in the C-terminal zinc finger domain. The ortholog to AFUB_013240 in the model mould *Aspergillus nidulans* is the repressor of sexual development *silG* (AN0709) with 56.3% identity and 67% similarity throughout the protein using EMBOSS Needle alignment (Martins et al., 2014). Thus, the molecular function and biological processes associated with AFUB_013240 remain to be clearly defined, but we herein name AFUB_013240 *silG* consistent with the gene name given in *A. nidulans*.

### SilG plays a role in biofilm development

Given the high levels of expression of *silG* in late-stage biofilms we hypothesized that SilG plays a role biofilm maturation or maintenance. To investigate this, we generated a *silG* null mutant strain by replacing the AFUB_013240 coding sequence with the hygromycin B resistance gene *hphB* in the strain CEA10. Given the previous low oxygen induction of *silG,* we first investigated if *silG* was essential for low oxygen fitness utilizing standard colony biofilm growth assays at 21% and 0.2% O_2_. Loss of *silG* did not impact radial growth or colony morphology under the tested low oxygen conditions compared to the wildtype strain, indicating that *silG* does not play a role in low oxygen colony biofilm growth or morphology **(Fig. S7)**.

We next assessed the impact of *silG* loss in the submerged biofilm model where we observed a striking time dependent increase *silG* transcript levels over time. Perhaps consistent with the transcript profile, the Δ*silG* strain has equivalent biomass to the wildtype and reconstituted strains at early timepoints (18- and 24-hours) but has a reduction in biomass in late-stage biofilms **(Fig. 8F).** This reduction is significant at the 30-hour timepoint, when *silG* expression is the highest. We additionally investigated the biomass of the Δ*silG* mutant at the 40-hour timepoint and observed a continued trend of reduced biofilm biomass (**Fig 8G**). Therefore, we conclude that SilG plays a role in the maintenance of mature *A. fumigatus* biofilms.

## DISCUSSION

Prior to our current study, the transcriptional landscape of the *A. fumigatus* sub-merged biofilm model was not described. Our understanding of the biology of this organism at a transcriptional level has come from shaking batch cultures and colony biofilm modes of growth (Gibbons et al., 2012; Hillmann et al., 2014; Kowalski et al., 2019; Losada et al., 2014; Muszkieta et al., 2013). However, neither of these modes of growth closely recapitulate the structure and environment of *A. fumigatus* growth *in vivo.* We have previously observed that our submerged culture biofilm model establishes similar fungal structure as *A. fumigatus* in murine model pulmonary lesions (Grahl et al., 2011; Kowalski et al., 2019, 2020). Therefore, we expect that analysis of a submerged biofilm will yield insights into pathogenesis and virulence relevant mechanisms. For example, support for this model’s relevance to discovering *in vivo* essential genes include the observation that the *srbA* null mutant fails to form a biofilm in this model and *in vivo,* and the herein observed impact of fermentation pathways on biofilm formation (Chung et al., 2014; Grahl et al., 2011; Kowalski et al., 2020; Willger et al., 2008). Moreover, the model was successful in identifying the alanine aminotransferase, *alaA,* as a mediator of echinocandin susceptibility in biofilms and in a murine model of IPA (Kerkaert et al., 2022).

Utilizing an RNA-Seq approach to define transcript abundance at selected time points of *A. fumigatus* biofilm development revealed temporally unique transcriptional profiles. A hallmark of these temporally distinct transcript profiles is dynamic changes in transcripts associated with metabolism. By looking directly at transcripts associated with primary metabolism, we find a shift in the transcript levels of genes associated with respiration (oxidative phosphorylation) and fermentation as the biofilm matures (**Fig. 5**). Although this is the first time this is described in *A. fumigatus* submerged biofilm model, the observation of metabolic shifts as the result of a biofilm mode of growth have been previously described in bacterial and other fungal biofilm systems (Delaney et al., 2023; Lu et al., 2019; Pisithkul et al., 2019; Rai et al., 2024). Similar to our model system here, in these systems there are metabolic shifts that occur during biofilm development and maturation. In our model fungal biofilm system, the shift is away from respiration to favor fermentation. The mechanism(s) driving this metabolic shift are unclear, but likely are the result of changing micro and macronutrient availability within the biofilm structure as development progresses. Importantly, considering the microenvironmental changes occurring within biofilms in this model raises a potential limitation; the primary carbon and nitrogen sources utilized by the fungus *in vivo* remain ill-defined. Thus, our data must be interpreted recognizing the use of glucose and nitrate as the primary carbon and nitrogen sources, respectively, for biofilm development in the studied model.

In support of the model’s *in vivo* relevance, a previous study identified the alcohol dehydrogenase, AlcC, as a mediator of fungal burden in a murine model of invasive pulmonary aspergillosis. The mechanism driving the reduction of fungal burden *in vivo* in the absence of *alcC* has remained ill-defined. Here in our analysis, we observed increases in transcripts associated with fermentation, including *alcC,* as biofilms developed and matured. Thus, one possibility for the reduction of fungal burden associated with loss of *alcC* is an inability to fully develop a mature biofilm in the murine lung environment. While loss of the alcohol dehydrogenase *alcC* did not significantly reduce biofilm biomass in the conditions we examined here, in the submerged model, additional loss of the *bdhA* gene involved in butanediol fermentation significantly reduced fungal biofilm biomass. Previously, 2,3-butanedione was detected in the headspace of *A. fumigatus* batch cultures in low oxygen conditions (Rees et al., 2017). It is possible that the multiple fermentation pathways induced in the *in vitro* submerged model are due to the large amount of glucose utilized as the primary carbon source in these experiments. Moreover, it is clear that *A. fumigatus* may produce other pyruvate derived fermentation products such as lactate. It will be important to further *dissect in* vivo relevant carbon and nitrogen sources and how they influence specific fungal fermentation pathways in established infections.

One approach to address the complex regulation of metabolism and fermentation and the associated metabolic products is to define the regulators of these pathways. We examined the transcript levels of 429 predicted transcription factors and observed significant abundance changes during biofilm development. Interestingly, we observed transcription factor encoding genes among our lowland transcripts at the 18-hour timepoint indicating the transcripts for many regulatory genes are down regulated at this early timepoint during development. From the heatmap for transcription factor transcript abundance (**Fig. 7A**), transcription factor transcripts are strongly regulated between the 12- and 18-hour and 24- and 30-hour timepoints. This data further highlights that there is an important change occurring between 12 and 18 hours and then again between 24 and 30 hours. These dramatic transcriptional changes could be the driver of principal component 2 (PC2) from the exploratory analysis where we found that samples separate on PC2 by early/late biofilms and middle stage biofilms **(Fig. 2B)**. What is driving these transcriptional changes is unclear. However, previous work from our group has shown oxygen gradients play a key role during biofilm development and are likely involved in driving the PC2 variability (Kowalski et al., 2020). Therefore, the transcriptional shifts are in part likely driven by oxygen gradient dynamics at a given timepoint. Moreover, as indicated by the genetic analysis of specific fermentation pathways, alternative carbon sources produced from glucose metabolism are likely also driving transcriptional changes. The reduction in *creA* transcript levels as the biofilm develops likely supports this hypothesis. Taken together, these data also highlight a general limitation in our approach using bulk-RNA-Seq of the biofilm model as it is likely substantial spatial heterogeneity exists within the biofilm. However, these data allow predictions to be made, and hypotheses tested with developing single cell and reporter gene approaches in the biofilm model.

We further tested the predictive power of the dataset by examining a specific, unstudied, predicted transcription factor that is regulated in part by oxygen levels, AFUB_013240, herein called *silG.* The transcript abundance for *silG* is significantly increased in hypoxic conditions in a SrbA independent manner (Chung et al., 2014; Kowalski et al., 2019; Losada et al., 2014). Interestingly, *silG* transcript abundance was also found to be significantly increased after 16-hours when co-cultured with host epithelial cells (Watkins et al., 2018). In the submerged biofilm model, *silG* transcript levels rise over the course of biofilm development, perhaps consistent with the decreasing levels of oxygen previously observed in mature biofilms. However, we cannot rule out that *silG* transcript levels respond to other changing culture conditions that are concomitant with reductions in oxygen levels. Interestingly, loss of *silG* did not impact colony growth or morphology on agar under low oxygen conditions. However, loss of *silG* in the submerged biofilm model resulted in a reduction of biofilm biomass. These results may suggest *silG* responds to a metabolic consequence of the fungal hypoxia response and is not a regulator of the hypoxia response per se. Importantly, SilG is required to maintain biofilm biomass in the submerged model, and arguably is the first filamentous fungal biofilm maintenance regulator identified. SilG has not been characterized fully in filamentous fungi to date, though it may be related to the proteosome regulator in yeast, RPN4. It is intriguing that *silG* has been associated with sexual reproduction in *A. nidulans,* which is promoted by oxygen limiting conditions. Future studies on the function of *silG* in *A. fumigatus* should include analysis of genes it regulates and its impact on pathogenicity and disease progress in murine models of aspergillosis.

## MATERIALS AND METHODS

### Strains and growth conditions

Mutant strains were made in the *Aspergillus fumigatus* CEA10 background, also called FGSC A1163; therefore, CEA10 was used as the wild-type (WT) strains as appropriate for each experiment. Strains were stored as conidia in 25% glycerol at −80°C and grown on 1% glucose minimal medium (GMM) at 37 °C (Shimizu & Keller, 2001). Conidia were collected using 0.01% Tween 80.

### Microscopy

Images of biofilms were taken using a Nikon spinning disk microscope fitted with a Yokogowa CSU-W1 spinning head and a 20x dry objective. Biofilms were grown for indicated times and stained with 25 µg/ml of calcofluor white 20 minutes prior to imaging. Three-dimensional image rendering was performed using BiofilmQ analysis software and Paraview visualization software (kitware) (Hartmann et al., 2021; Kowalski et al., 2019).

### RNA extraction from biofilm tissue

Submerged liquid cultures were seeded in GMM with 10^5^ spores per mL in 100mm petri dishes. Biomass was collected via filtration through miracloth and snap frozen in liquid nitrogen. Frozen biomass was bead beat in 200uL of TRI Reagent (Invitrogen) with 2.3-mm silica beads. Homogenate was brought to a final volume of 1 mL with TRI Reagent and RNA was extracted by following the manufacturer’s protocol.

### RNA-sequencing

RNA for RNA-seq was quantified by qubit (Thermo Fisher Scientific) and integrity measured on a fragment analyzer (Agilent). Samples with RIN ≥ 7 underwent library preparation with the mRNA HyperPrep kit (Kapa Bioscience) using 200ng RNA as input following manufacturer’s instructions. Libraries were pooled for sequencing on a NextSeq500 instrument (Illumina), targeting 10M, single end 75bp reads/sample.

### RNA-sequencing analysis

Read alignment, normalization, and differential gene expression analysis. Raw read quality was initially assessed using fastQC (Babraham Bioinformatics group) and multiQC software (Ewels et al., 2016). Reads were trimmed using Cutadapt software using a quality score cutoff of 20 and again assessed for quality using fastQC and multiQC (Martin, 2011). Alignments were performed using Star alignment software with the *A. fumigatus* CEA10 reference genome version FungiDB-52 and general feature format file from same version (Dobin et al., 2013). Counts per gene were compiled using htseq (Anders et al., 2015; Dobin et al., 2013).

For the exploratory analysis the package DESeq2 was used to generate vst normalized values for use in clustering and principal component analysis (Love et al., 2014). The package edgeR was used to normalize read data for use in differential expression analysis using the TMM method (Robinson et al., 2010). Low abundant transcripts were filtered using the fitlerByExpr function and normalization was performed using the CalcNormFactors with using trimmed mean of the M-values. Limma-voom methodology was used to perform a differential expression analysis using linear model fit with Empirical Bayes moderated t-statistics and multiple comparisons between all pairwise timepoint combinations (Ritchie et al., 2015). Scaled CPM values are used for heatmap representation using the R scale function which works by subtracting values from the mean CPM for each gene and dividing by the standard deviation for the same gene. Heatmaps were generated using the ComplexHeatmap package (Gu, 2022). Upset plots were generated using the UpSetR package (Conway et al., 2017; Lex et al., 2014).

### Strain construction

The Δ*silG* null mutant was generated by replacing the *silG* open reading frame (AFUB_013240) with the dominant selection marker *hphB* in the CEA10 background utilizing a CRISPR/Cas9 mediated transformation into protoplasts as previously described (Al Abdallah et al., 2017; Kerkaert et al., 2022). The repair construct with the *hphB* resistance marker was amplified using RAC7880 and 7881. This amplified repair was transformed into CEA10 protoplasts using CRISPR/Cas9-mediated targeting to the *silG* locus. Mutants were selected for growth on osmotically stabilized media containing hygromycin. Loss of *silG* was confirmed by Southern blot analysis using DIG-labelled probe generated by primers RAC8065 and RAC8066 against 30ug of WT and Δ*silG* gDNA digested using EcoRV-HF (NEB) restriction digest enzyme in CutSmart Buffer (NEB) for 16-hrs at 37°C. Digested DNA was run on 1% agarose gel for 2-hrs at 80V, transferred to PVDF membrane using vacuum blotter, and then probed overnight with DIG-labelled DNA probe (Roche Diagnostics, Mannheim, Germany). Antibody detection was done according to manufacturer protocol (Roche). Southern blot was visualized using chemiluminescent gel imaging system, DIG-labeled Molecular Weight Marker VII was used for DNA ladder (Roche).

Reconstitution of the *AFUB_013240* gene was performed by PCR amplification of the *AFUB_013240* locus from wildtype gDNA from ∼1000 bp upstream of the start codon to ∼1000 bp downstream of the stop codon using primers RAC8178 and RAC8179 with homology to a backbone plasmid containing the pyrithiamine resistance marker *ptrA*. The backbone plasmid was amplified using primers RAC8180 and RAC 8181. The resulting PCR products were assembled using NEBuilder Hi-Fi DNA assembly kit (NEB). A repair construct was amplified from this resulting plasmid using primers that contained homology to the *aft4* safe-haven locus using primers RAC8184 and RAC8185 (Furukawa et al., 2022). The repair construct was transformed into Δ*silG* protoplasts using CRISPR/Cas9-mediated targeting to the *aft4* safe-haven site. Mutants were selected for growth on osmotically stabilized media containing pyrithiamine. Reconstitution of gene expression was validated by RT-qPCR on RNA extracted from 24-hr WT, Δ*silG,* and *silG*^recon^ biofilms using the primers RAC8464 and RAC8465.

The Δ*bdhA* null mutant was generated by replacing the *bdhA* open reading frame (AFUB_031610) with the *pyrG* selection marker in the uridine/uracil auxotrophic CEA17 background. RAC4242 and RAC4239 were used to amplify a ∼1.8kb region 5’ to the *bdhA* open reading frame and RAC4240 and RAC4243 were used to amplify a ∼850bp region 3’ to the open reading frame. The *pyrG* gene along with its promoter and terminator were amplified using RAC2055 and RAC2056. The two flanks and the *pyrG* marker were combined via overlap PCR and the resulting product was used to transform CEA17 protoplasts as previously described (Kerkaert et al 2022). Mutants were selected for growth on osmotically stabilized minimal media lacking uridine/uracil supplementation (GMM plus 1.2M sorbitol). Similarly to generate the Δ*bdhA*Δ*alcC* double mutant, the *bdhA* open reading frame was replaced with a pyrithiamine resistance selection marker in a previously published Δ*alcC* strain (Grahl et al., 2011). RAC2055 and RAC2056 were used to amplify the pyrithiamine selection marker and the resulting product was combined with the two previously mentioned ∼1.8kb and ∼850bp flanks via overlap PCR.Mutants were selected on osmotically stabilized minimal media containing pyrithiamine. Both *bdhA* null strains were single spored and checked for correct locus integration via PCR and southern blotting.

The *bdhA* reconstitution construct was generated by amplifying the *bdhA* ORF ± 1000bp up- and down-stream from *bdhA* using RAC8921 and RAC8922 and the plasmid backbone containing the *hphA* resistance cassette with RAC8923 and RAC8924. The bdhA cassette was integrated into the plasmid backbone by HiFi DNA assembly using NEBuilder (NEB) resulting in the pbdhA-hphA plasmid. The reconstitution repair construct was PCR amplified from pbdhA-hphA using RAC8964 and RAC8965 which contain homology to the *aft4* safe-haven locus and transformed into Δ*bdhA* protoplasts as described above. Expression levels were analyzed using RT-qPCR on RNA extracted from 24-hr biofilms using the primers RAC8955 and RAC8957.

All strains were single spored and checked for correct integration via PCR and Southern blotting. Primer sequences used are available in Table S5. All strains generated in this study are available upon request from the corresponding author.

### RT-qPCR

A sample of 5 μg of RNA was DNase treated with the TURBO DNA-free kit (Invitrogen) according to the manufacturer’s protocol. A sample of 500 ng of DNase-treated RNA was run on an agarose gel to ensure RNA integrity, and 500 ng of DNase-treated RNA was used for cDNA synthesis as previously described (Beattie et al., 2017). The RT-qPCR data were collected on a CFX Connect real-time PCR detection system (Bio-Rad) with CFX Maestro Software (Bio-Rad). Gene expression was normalized to *actA* and *tefA* expression for all experiments. Primers used are available in the primer table (Table S5).

### Growth assays

Agar plates were inoculated with 10^3^ conidia and incubated for 72 h at 37°C, 5% CO_2_ in either ambient oxygen or in a chamber that maintained oxygen at a concentration of 0.2% using an InvivO_2_ 400 Workstation (Ruskinn Baker).

Biofilm biomass cultures were inoculated with 2 mL of 10^5^ conidia/mL in GMM and grown in 6-well tissue culture plates for indicated time (12, 18, 24, or 30-hrs) at 37°C, 5% CO_2_. Supernatants were removed, and biofilms were harvested via transfer to conical tubes and centrifugation. Biomass was washed 2× with double-distilled water, frozen at −80°C, lyophilized, and dry weight measured. For the 40-hour timepoints 4ml of 5×10^4^ conidia/ml was used to prevent the formation of growth at the air liquid interface. For 40hr experiments, parallel 18hr biofilms were also grown in 4ml of 5×10^4^ conidia/ml to keep comparisons consistent between two time points.

### Acetoin Quantification with modified Voges-Proskauer assay

Strains were grown in GMM with 5×107 conidial in 100mL, 200rpm, 37°C, and 5% CO2, for 24-hours. Fungal biomass was collected and swapped into 100mLs of fresh GMM and cultured for 48 hours in hypoxia, 1% oxygen (200rpm, 37°C, 5% CO2). Mycelia were harvested, rinsed with distilled water, frozen and lyophilized. Culture supernatants were collected. Acetoin standards (0-1.0mM) as well as experimental samples were incubated with 0.5% creatine, 5% alpha-naphthol, and 40% potassium hydroxide for 15 minutes. Absorbance was measured at 560nm and mM concentration of acetoin calculated based on standard curve (Ljutov, 1963; Rinehart et al., 2018).

### Statistical Analysis

All statistical analyses were performed in GraphPad Prism 10. Error bars indicate standard deviation around the mean.

### Data availability

The data discussed in this publication have been deposited in NCBI’s Gene Expression Omnibus (Edgar *et al*., 2002) and are accessible through GEO Series accession number GSE298422 (https://www.ncbi.nlm.nih.gov/geo/query/acc.cgi?acc= GSE298422).

## ACKNOWLEDGEMENTS

This work was funded by NIH/NIAID grants R01AI130128 and R01AI146121 (RAC). KAM was also supported by an NIH/NHBLI T32HL134598 grant (O’Toole, George). JDK and KGQ were also supported by an NIH/NIAID T32AI007519 (Hogan, Deborah). Core facility support provided by NIH grant P20-GM113132 to the Dartmouth BioMT COBRE. Additional support was provided by the Cystic Fibrosis Foundation Research Development Program (STANTO19R0) and NIH/NIDDK P30-DK117469 (Dartmouth Cystic Fibrosis Research Center). The authors would also like to thank Ann Lavanway in the Dartmouth Microscopy core for assistance with imaging.

---

**Figure S1:**
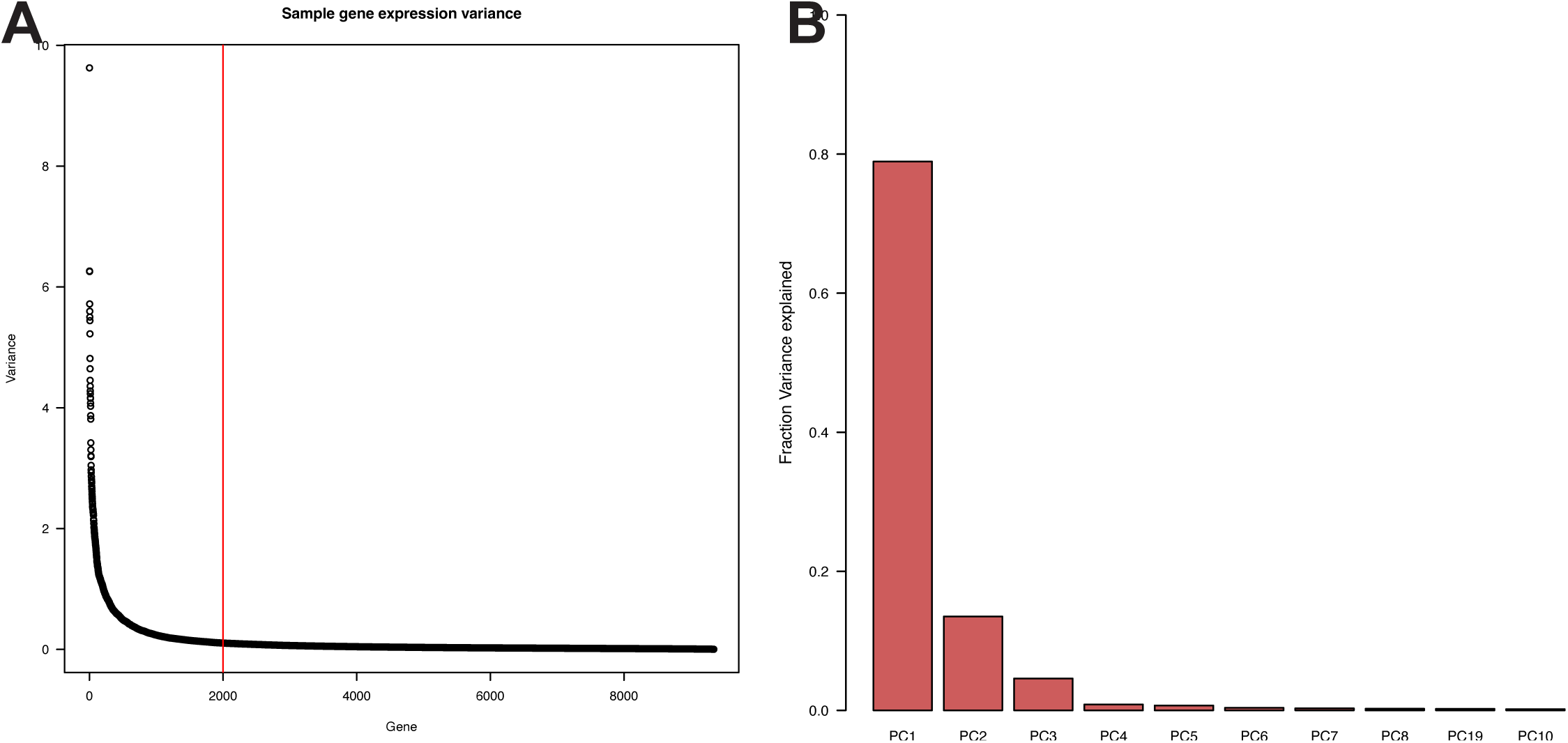
**A)** Genes were ordered by variance to identify a threshold of 2000 genes to use for PCA and variable gene expression heatmap. **B**) Principal component analysis reveals the first 2 principal components captures the majority of variation.

**Figure S2:**
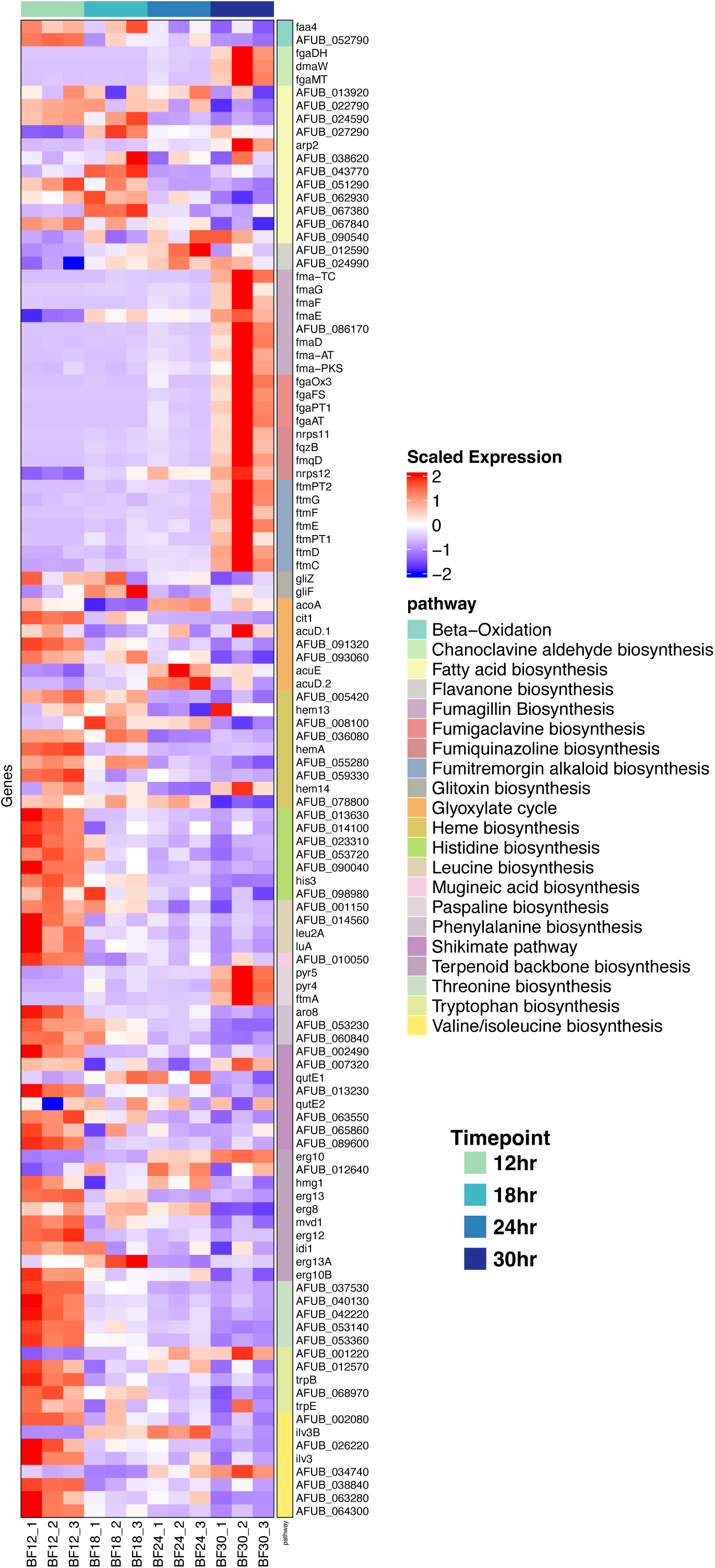
A heatmap of secondary metabolism and related pathways. Pathways were identified from the KEGG database. Scaled CPM values are shown.

**Figure S3:**
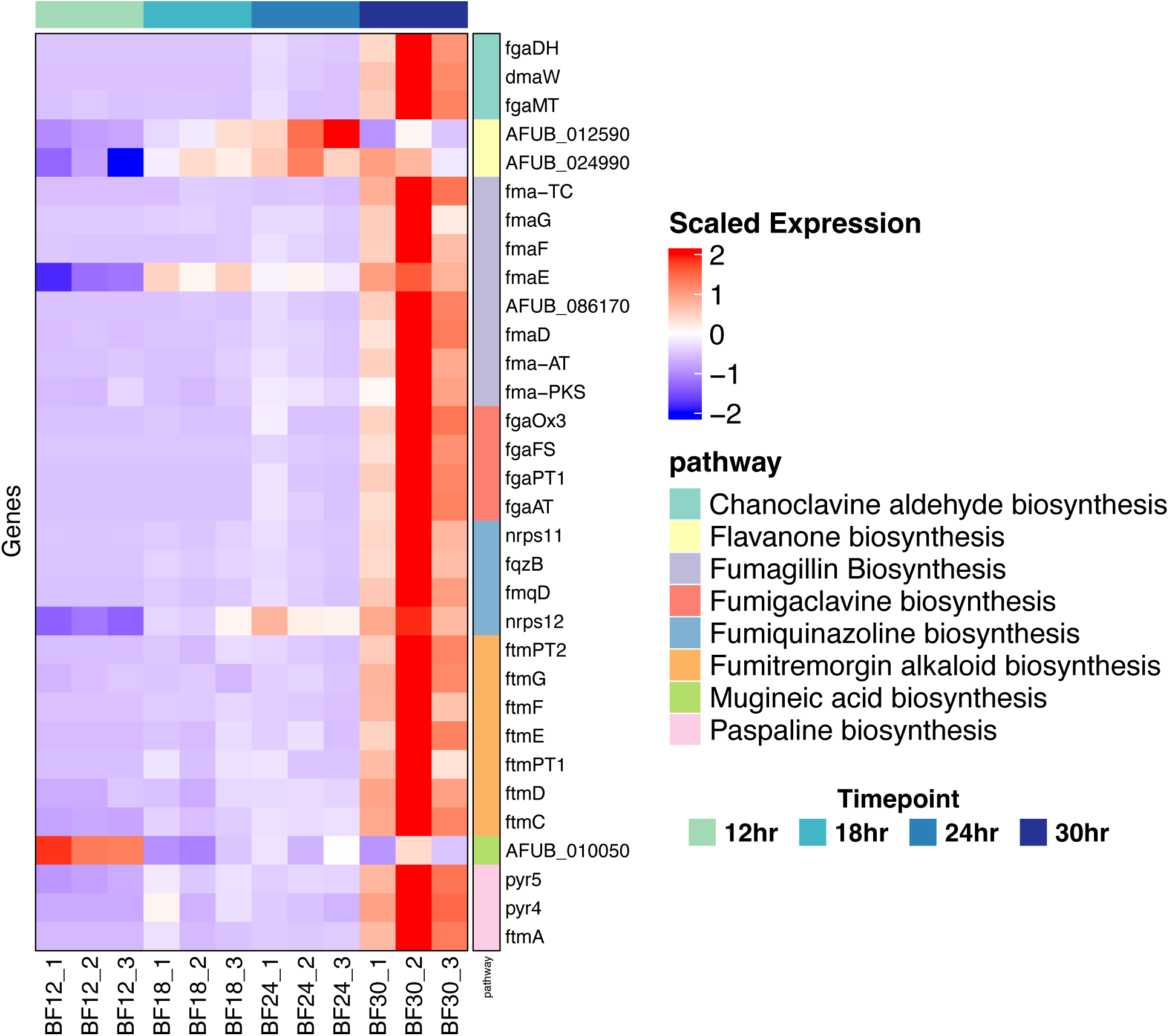
A heatmap of secondary metabolite biosynthetic gene clusters transcript abundance. Several of these gene clusters are up regulated in late stages of biofilm development. Scaled CPM values are shown.

**Figure S4.**
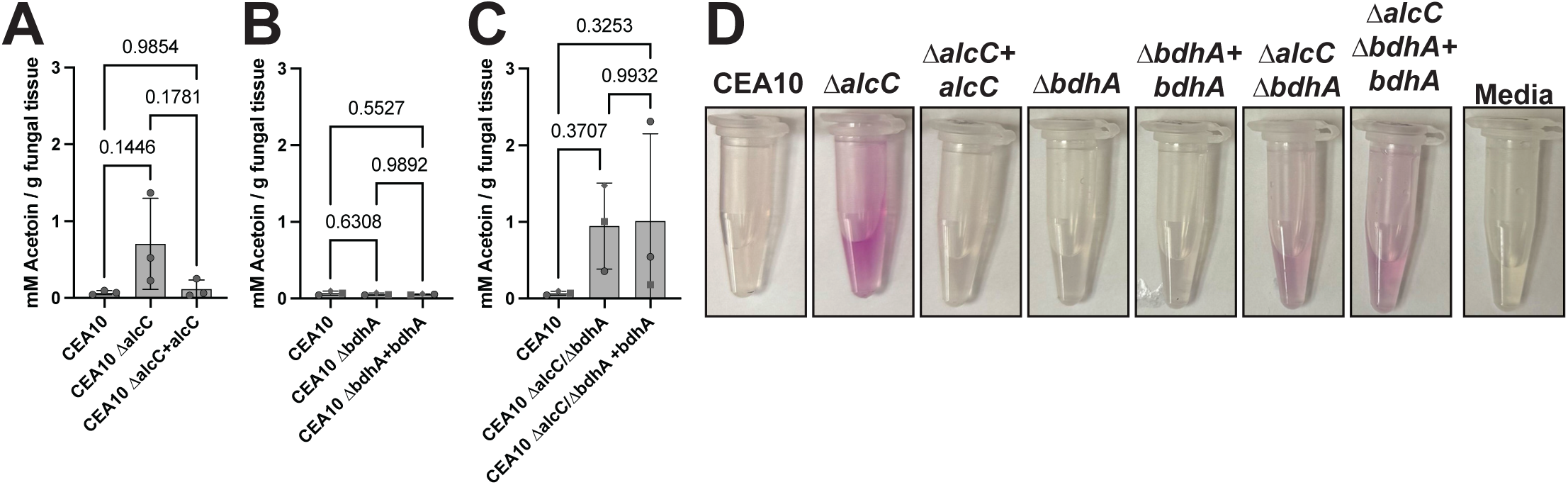
Acetoin accumulation in culture supernatants of low oxygen cultures. Acetoin levels were quantified in culture supernatants from shaking cultures that were grown for 24 hours in atmospheric oxygen supplemented with 5% CO_2_ switched into 0.2% O_2_ supplemented with 5% CO_2_ conditions for 48 additional hours. **A)** Quantification of acetoin in culture supernatants using a Voges-Proskauer assay. **B**) representative images of colorimetric output from acetoin detection analysis. Statistics are a one-way ANOVA with a Tukey’s multiple comparison test.

**Figure S5:**
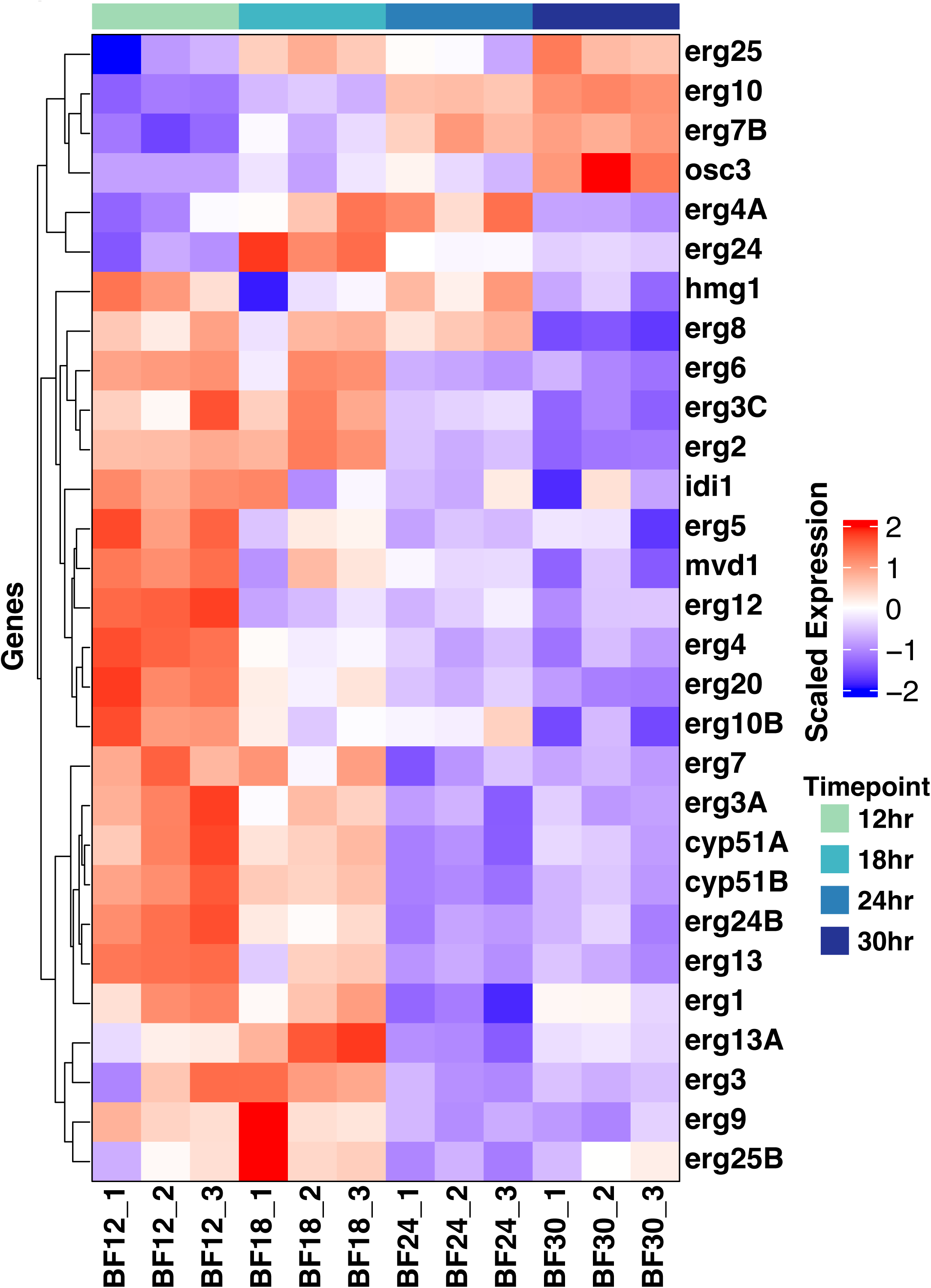
Heatmap of the ergosterol pathway with hierarchical clustering. Scaled CPM values are shown.

**Figure S6:**
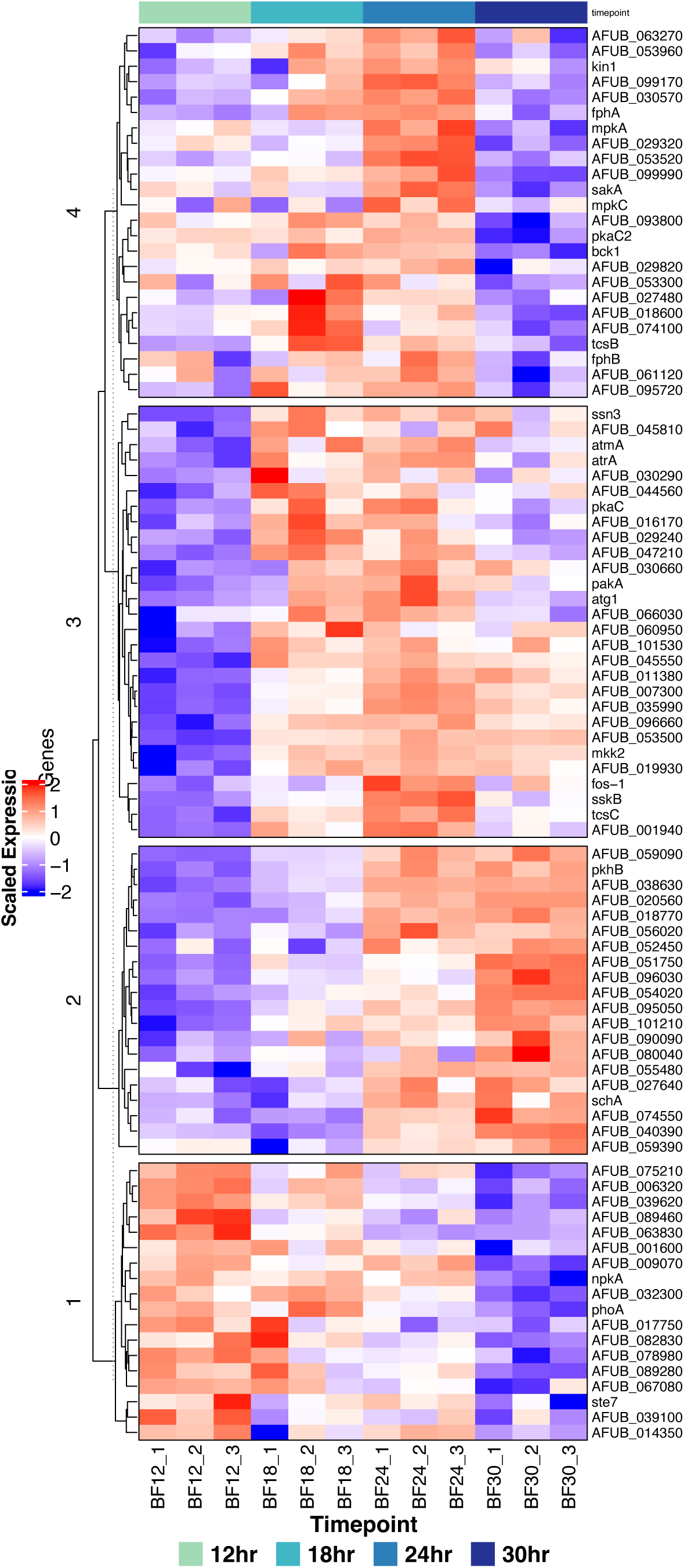
Heatmap of kinase transcript abundance with hierarchical clustering. Scaled CPM values are shown.

**Figure S7.**
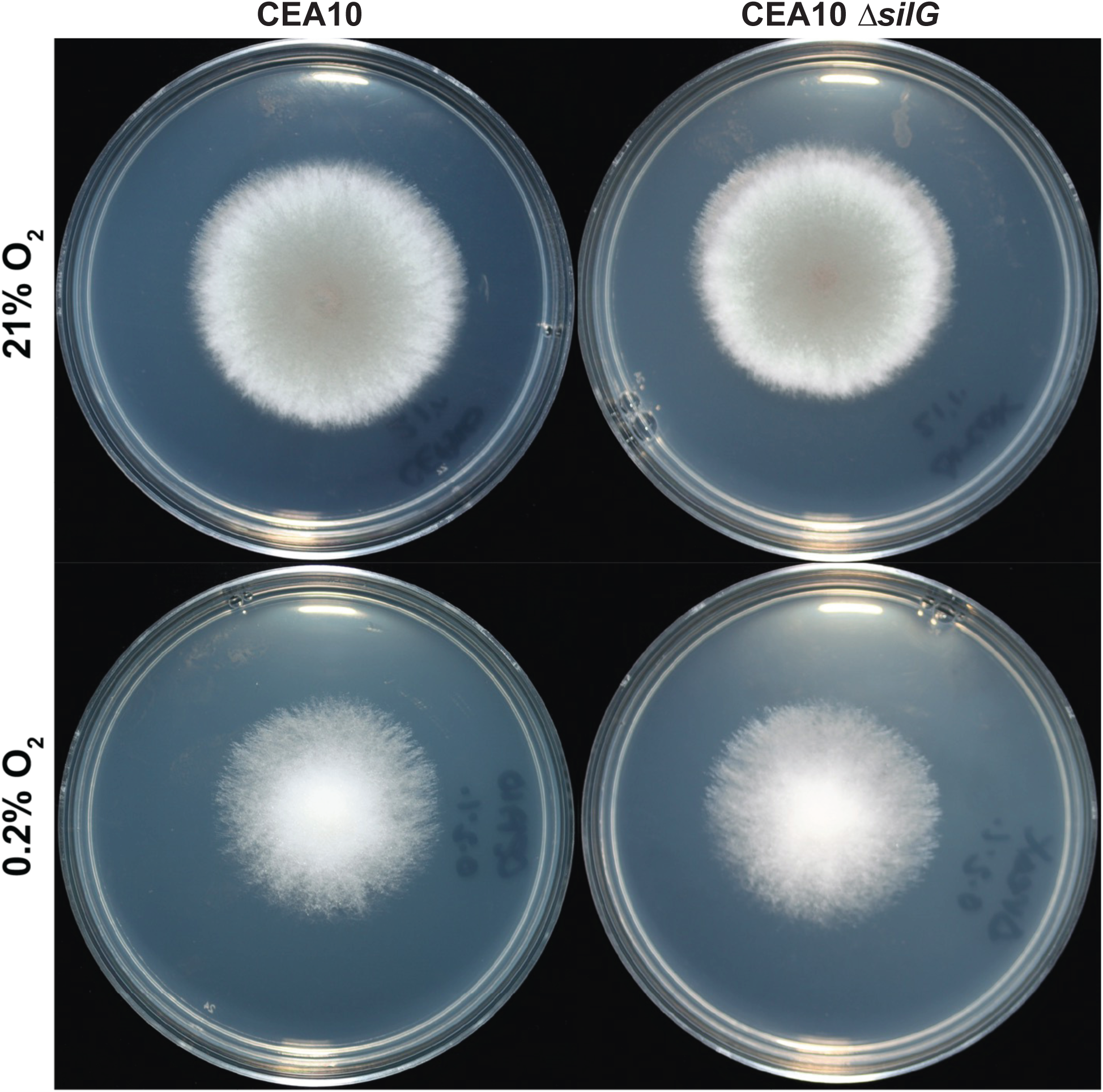
*AFUB_013240* is dispensable for colony biofilm growth in hypoxia. Representative images of colony biofilm morphology of Δ*AFUB_013240* growth on solid glucose minimal media at 21% and 0.2% O_2_ compared to wildtype CEA10.

**Table S1.** Table of top 2000 most variable genes with DESeq2 vst normalized values.

**Table S2.** Table of log2FC values and adjusted p-values from pairwise differential expression analysis.

**Table S3.** Genes and log_2_ fold-change information for genes in peak, plateau, lowland, and valley clusters at each timepoint.

**Table S4.** FunCat category enrichment analysis for dynamic transcripts. Only significantly enriched categories are included.

**Table S5.** DNA oligonucleotides used in this study.

## Notes

Conflict of Interest Statement: The authors have declared that no conflict of interest exists.

